# Association of Oxytetracycline and Florfenicol affects zebrafish larvae behavioral repertoire

**DOI:** 10.1101/2023.04.20.537675

**Authors:** Darlan Gusso, Marilia Oberto da Silva Gobbo, Gabriel Rübensam, Carla Denise Bonan

## Abstract

The fish farms are increasingly growing and, consequently, the use of antibiotics in aquaculture. Oxytetracycline (OTC) and Florfenicol (FF) are the most common antibiotics applied in the fish production sector and livestock farming. The elucidation of the effects of the antibiotics is essential to control their use and their physiological and pharmacological implications. Here we studied the behavioral effects of the 96 h-exposure to OTC (2, 10, 20, and 100 mg/L), FF (0.01, 0.05, 2 and 10 mg/L) or OTC (10 mg/L) + FF (10 mg/L) in zebrafish larvae. We observed that the covered distance and the movement increased in animals exposed to OTC + FF when compared to control. In addition, fish entered the center of the plate test more often and stayed there longer. The turn angle was reduced at OTC + FF. We also observed that the optomotor response was compromised by 10 and 20 mg/L OTC and to OTC + FF. Our data demonstrated that there is an increase in the number of entries in the center area and time spent in center area for FF- and OTC + FF-treated groups. These data showed that the antibiotics promoted a reduction of anxiety-like behavior allowing larvae to explore more the novel environment as well as a detrimental performance for the optomotor response.

**Highlights:**

- Florfenicol (FF) did not alter exploratory and anxiety-like behavior in zebrafish larvae.
- Oxytetracycline (OTC) did not alter exploratory behavior, but there was an increase in the time spent in the center area
- OTC plus FF increased distance and movement in zebrafish larvae.
- OTC plus FF reduced the anxiety-like behavior in zebrafish larvae.
- Optomotor behavior was compromised by treatments with OTC or OTC + FF.

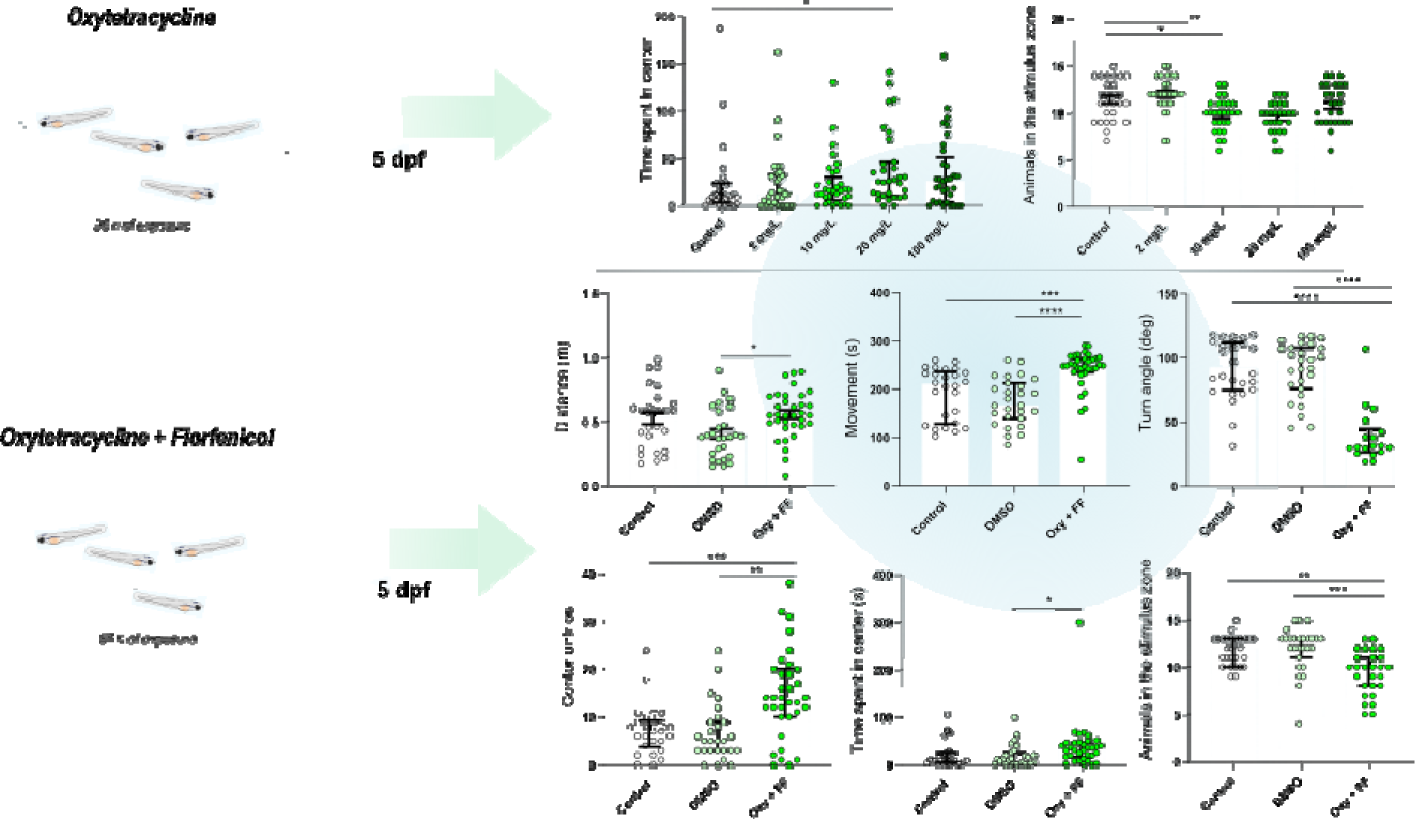

## Introduction

Aquaculture industries play an important role in supplying quality protein for human consumption. Fish consumption has grown rapidly reaching a range of 20.5 kg per capita in 2018. This growth follows an annual world production of 179 million tons (FAO, 2020). Global trends point to an increase in fish production to 204 million tons until 2030 following a population growth of 8.5 billion people (FAO, 2020; ONU, 2019; Schar et al., 2020). The aquaculture industry offers enormous potential to respond to the demand for food associated with global population growth. Although since the 1980s fish capture has been losing ground to conventional breeding high densities systems (FAO, 2020), this scenario has shifted since 2001 when global aquaculture achieved an average growth of 5.8 that was mainly attributed to the increase in demand for protein of animal origin (Schar et al., 2020).

Globally, fish farming has been improving its environmental performance imposed by government regulation and responding to market demand for protein from biosustainable systems (Naylor et al., 2021). Following those improvements, the implementation of controlled and intensive fish production systems have been essential (FAO, 2020; Naylor et al., 2021; Schar et al., 2020).

As one of the world’s fastest-growing food production sectors, aquaculture is experiencing an escalation in the use of antibiotics to keep farmed fish free of disease. As a result of that, an increase in the discharge of antibiotics into the environment has been alarmingly observed (Wernli et al., 2020; Yu et al., 2020). The use of antibiotics in aquaculture per kilogram of biomass (164.8 mg kg ^- 1^) exceeds its medicinal use in humans (92.2 mg kg ^- 1^) as well as in terrestrial animal farming (140 mg kg ^- 1^) (Schar et al., 2020). Despite the high range of antibiotics in fish production, each country has independently been regulating its antibiotics use. The largest fish producer is China and it accounts for 35% of all fish produced in 2018 (FAO, 2020). Among several molecules approved for fish production, Florfenicol and Oxytetracycline in almost all countries have been used. Both compounds are most applicable for tilapia and salmon.

Florfenicol (FF) is a broad-spectrum antibiotic applied to treat aquaculture infections and is widely and exclusively for veterinary medicine use (Saito-Shida et al., 2019). This antimicrobial is effective against a range of infectious diseases. Currently, its use is indicated to treat or prevent a salmon infection caused by *Piscirickettsia salmonis* (San Martín et al., 2019), a coldwater disease associated with *Flavobacterium psychrophilum* and furunculosis associated with *Aeromonas salmonicida*. It is also used to prevent or treat enteric septicemia caused by *Edwardsiella ictaluri*, columnaris associated with *Flavobacterium columnare,* and streptococcal septicemia associated with *Streptococcus iniae* and *agalactiae* in a range of freshwater fish species such as catfish, tilapia, and finfish (FDA - Food and Drug Administration, 2021).

Oxytetracycline (OTC) is a broad-spectrum antibiotic applied to veterinary and also human use (Almeida et al., 2019; Rok et al., 2017). OTC dihydrate and hydrochloride present distinct administration ways. OTC dihydrate is administered feed-mixed and OTC hydrochloride is used in the bath treatment. The most common applications of OTC dihydrate can prevent or treat fish against *Hemophilus piscium* that causes ulcer disease, *Aeromonas salmonicida* which causes furunculosis, *Aeromonas liquefaciens* that causes bacterial hemorrhagic septicemia, *Aerococcus viridans* that causes gaffkemia, *Flavobacterium psychrophilum*, *Flavobacterium columnare,* and *Pseudomonas spp.* in a wide variety of fish, salmon, catfish, and tilapia. OTC hydrochloride is applied to prevent or treat finfish/fry and fingerling during a critical survival period from before hatching to juvenile stages.

Almost all countries use antibiotics for fish production with a variation of quantity and active ingredient (EMA - European Medicines Agency, 2020; FDA - Food and Drug Administration, 2021; Henriksson et al., 2018; Mo et al., 2017). Thus, it is important to evaluate that antibiotic use and its range are constantly increasing around the world. This situation might have serious consequences related to compensatory antibiotic resistance (Laxminarayan et al., 2013; Schar et al., 2020). As a result, the variation of antibiotics used in fish treatment has been demonstrated to have a diversity of effects ranging from behavior, including locomotor activity, defense against predators, and neurological impairment to microbiome dysbiosis (Li et al., 2020; Navarrete et al., 2008).

Several theories for the development of abnormalities caused by antibiotics are being discussed. In this sense, it is essential to study the role of antibiotics using zebrafish, a useful animal model to understand the mechanisms behind these disorders. In recent decades, zebrafish have been used for the development of studies in various areas, such as pharmacology (Idalencio et al., 2015; Petersen et al., 2021) toxicology (Bridi et al., 2017; Gusso et al., 2020), and endocrinology (Gaspary et al., 2018; Giacomini et al., 2015). This model has gained space for studies in neurosciences, contributing to a better understanding of brain function and dysfunctions and also to pharmacological modulation (Grossman et al., 2011; Müller et al., 2020). In addition to presenting sensitivity to drugs, the zebrafish model has a wide behavioral repertoire well described (Gerlai, 2016, 2003; Miller and Gerlai, 2008). The impacts caused by the constant fish exposure to antibiotics, especially related to neurotoxicity, are still poorly studied. Therefore, as this situation represents a challenge to global aquaculture, it became essential to examine and analyze the role of antibiotics on behavior using zebrafish as an animal model (Almeida et al., 2019; Kayani et al., 2021; Petersen et al., 2021).

Considering the use of the zebrafish as an animal model for environmental and translational studies and the common use of antibiotics in fish farms, this study aimed to evaluate OTC, FF, and FF + OTC effects on zebrafish larvae behavior.

## 2. Materials and methods

### 2.1 Study strategy

Zebrafish represent a sensitive organism to evaluate drug exposure through behavior analysis. Here we described the behavioral effects caused by OTC or FF alone, and both OTC + FF compounds. The antibiotics combination is a routine practice since they are administered to fish in the larval stage during the growth process. Also, both compounds were used in aquaculture production (Harnisz et al., 2015; Monteiro et al., 2016). We tested 10 mg/L in antibiotics combination, considering the previous results obtained by this concentration (Gusso et al., 2021).

### 2.2 Zebrafish maintenance

The larvae were obtained from a core zebrafish facility following established practices. Zebrafish larvae (*Danio rerio*) wild type (AB strain) were used. Each plate (9 x 9 cm) sustained 20 larvae until 4 dpf with 30 mL water. For the experiments, larvae were caught randomly. The progenitors are maintained in an integrated aquarium system (Zebtec, Tecniplast®, Italy). The Zebtec contains reverse osmosis filtered water at the recommended temperature (28 °C ± 2 °C), pH (7.0 - 7.5), conductivity (300–700 μS), hardness (80–300 mg/L), ammonia, nitrite, nitrate, and chloride levels for this species. In the night before mating (17:00), animals were transferred to a breeding tank in which males and females were separated by a transparent partition. This partition was removed (08:00 am) and after 30 min, fertilized embryos were cleaned and placed in Petri dishes with system water. The photoperiod was 14 h light: 10 h dark. Animal’s diet was based on feeding with commercial flake and artemia (Westerfield, 2007). A greenhouse B.O.D (Biochemical Oxygen Demand) with standard temperature and photoperiod was used for larvae maintenance. All protocols were approved by the Institutional Animal Care Committee (CEUA: 8950, 2018) and followed the “Principles of Laboratory Animal Care” from the National Institutes of Health (NIH). This study was registered in the Sistema Nacional de Gestao do Patrimonio Genetico e Conhecimento Tradicional Associado-SISGEN (Protocol No. A3B073D).

### 2.3 Antibiotics exposure

The exposure to OTC, FF, and OTC + FF was carried out as follows. Zebrafish larvae at 4 dpf were exposed to OTC (2, 10, 20, and 100 mg/L; CAS number 79–57-2), FF (0.01, 0.05, 2, and 10 mg/L; CAS number 73231-34-2) or OTC+ FF (10 mg/L + 10 mg/L) for 96 h. The FF was diluted in Dimethyl sulfoxide (DMSO; CAS number 67- 68-5) at a final concentration of less than 0.1% (Gusso et al., 2020; Nery et al., 2014). Twenty larvae were kept in Petri dishes with 30 mL water for 4 dpf. The treatments were completely changed every day. Each exposure was conducted in triplicate before each behavioral test.

### 2.4 Mass Spectrometry (LC-MS/MS) for OTC and FF analysis

All collected samples were stored at −20 °C before analysis. The concentrations of OTC and FF solutions were quantified by liquid chromatography coupled to mass spectrometry (LC-MS/MS), using a Xevo TQ-S microsystem (Waters, Milford, MA, USA) in the treated water samples. Chromatographic separation was performed in a ZORBAX RRHT Extend-C18 (2.1 x 50 mm, 1.8 µm, Agilent, Paolo Alto, USA) column with a mobile phase consisting of (A) 0,1% of formic acid 2 mM ammonium acetate and (B) methanol with 0.1% formic acid and 2 mM of ammonium acetate, in gradient mode, at a flow rate of 0.4 mL/min. The gradient initiated with 5% of mobile phase B, going to 80% of B after 1.50 min, remaining in this proportion for 3 min. After the elution of the analytes, the initial condition was restored and the system was re-equilibrated for 0.5 min. The temperature of the chromatographic column was maintained at 50 °C and the injection volume was 5 µL. The analytes were ionized in an electrospray source (ESI) operated in negative and positive modes, utilizing nitrogen desolvation gas with a flux of 10 L/min, cone of 11 L/min, and capillary of 3500V, with the temperature at 550 °C. The collision gas was argonium, and the capillary of the ESI was adjusted to 1.0 KV. The spectrometer was operated with multiple reaction monitoring (MRM) mode with the following transitions (m/z): oxytetracycline with quantification ion 461>426 and confirmation ion of 461>381, and florfenicol with quantification ion of 356>336 and confirmation ion of 356>185. Before analysis, all samples were thawed at ambient temperature, agitated at 300 rpm for 10 min in an orbital mixer, and diluted with 5% of mobile phase B depending on the concentration. Standards were dissolved in methanol (1 mg/mL) and diluted with 5% mobile phase B to obtain calibration curves in the range of 5 to 200 ng/mL for both analytes. Quantification was performed by external standardization and the results were expressed in mg/L after dilution correction. The experiments were performed in triplicate (n=3 for each concentration tested)

### 2.5 Exploratory behavior

The locomotor parameters were assessed using 5 dpf larvae. The larvae were individually placed in 24 well cell culture plates filled with 3 mL of system water at 28 ± 2°C. They were acclimated for 1 min and tracked for 5 minutes. The exploratory behavior was registered by a tracking device (Noldus Information Technology, Wageningen, Netherlands). Zebrafish exploratory behavior test was assessed by distance covered (m), movement (s), acceleration (cm/s²), latency to first center entry (s), center entries, and time spent in center (s). All data were assessed using EthoVision XT 10.0 Software. A specific parameter movement was previously calibrated to consider the period during which the zebrafish exceeded the start velocity (0.06 cm/s) and remained moving until reaching the stop velocity (0.01 cm/s) (Colwill and Creton, 2011).

### 2.6 Aversive behavior

Through a red ball, a visual aversive stimulus was assessed. Larvae were placed in a 6-well plate (5 larvae per well - 3mL water) over an LCD monitor for ball avoidance behavior from a visual stimulus (a 1.35 cm diameter red bouncing ball) for a 5-min session following 1 min of acclimation (Nabinger et al., 2018). The sessions were video-recorded and analyzed with the naked eye by two different investigators. A red bouncing ball traveled from left to right over a straight 2 cm trajectory on the top half of the well plate area (stimulus area), which animals avoided by swimming to the other (non-stimulus) half of the well. After acclimation time, each video was stopped every 30 seconds to quantify the number of larvae in the non-stimulus and stimulus zones. The procedure considered the time percent at which each larva stayed in a non-stimulus area. The number of larvae in the non-stimulus area for all 5-min sessions indicated deficits in the avoidance response. The results were demonstrated as % of larvae in non-stimulus area.

### 2.7 Optomotor response

Optomotor responses were assessed through non-aversive visual environmental cues. At 5 dpf larvae were placed in Petri dishes (15 larvae per dish) over a LCD monitor and were exposed to moving white and black stripes (24.5 cm wide and 1.5 cm high) (Colwill and Creton, 2011). The stripes moving direction alternated between left and right every 1 min, with 5 s intervals in which they faded before reappearing. For the data analysis, the dish area was virtually divided into two zones (left and right) and the number of animals in the stimulus zone (the region where the pattern has moved) were counted during the 5-s white screen interval. Optomotor behavior follows the ability of larvae to respond to visual stimuli.

### 2.8 Statistical analysis

Data were expressed as mean ± standard error of the mean (S.E.M). A significance level of p < 0.05 was considered. Normality and distribution were evaluated by the Shapiro-Wilk test. Normal data were analyzed using one-way analysis of variance (ANOVA) followed by the Tukey test as a *post hoc*. Non-normal data were analyzed by the Kruskal-Wallis following Dunn’s multiple comparisons test. GraphPad Prism 8 (La Jolla, CA, USA) software was used for statistical analysis.

## 3. Results

### 3.1 Quantification of OTC and FF by LC-MS/MS

The OTC and FF concentrations in treated water were quantified through LC-MS/MS. The analytical concentration was close to nominal concentrations (Table 1).

**Table 1.** OTC and FF quantification in water using liquid chromatography coupled to tandem mass spectrometry (LC-MS/MS).

| Compound | Nominal concentration mg/L | Analytic concentration mg/L | Standard deviation |
| --- | --- | --- | --- |
| Oxytetracycline | 2 | 2.168 | 0.245 |
| Oxytetracycline | 10 | 12.493 | 1.347 |
| Oxytetracycline | 20 | 21.024 | 3.716 |
| Oxytetracycline | 100 | 104.537 | 6.436 |
| Florfenicol | 0.01 | 0.007 | 0.001 |
| Florfenicol | 0.05 | 0.057 | 0.003 |
| Florfenicol | 2 | 2.247 | 0.226 |
| Florfenicol | 10 | 11.612 | 3.075 |
| OTC + FF | 10 + 10 | 10.918 + 13.913 | 0.37 / 1.18 |

The data are expressed as mean ± standard deviation (SD) (n=3 for each concentration)

### 3.1 Locomotor activity

For OTC-treated animals, there were no changes in distance covered (F_(4, 155)_ = 1.474, p = 0.2128; Fig. 1A), movement (H = 5.700, p = 0.2227; Fig. 1D), and acceleration (F_(4, 163)_ = 0.8621, p = 0.4881 Fig. 1G). In FF-treated animals, we did not observe any changes in distance covered (H = 7.892, p = 0.1623; Fig. 1B), movement (F_(5, 155)_ = 2.227, p = 0.0543; Fig. 1E), and acceleration (H = 13.71, p = 0.0176; Fig. 1H). The OTC + FF did not present differences in distance covered (F_(2, 84)_ = 4.434, p = 0.8489; Fig. 1C) or acceleration (H = 0.9836, p = 0.6115; Fig. 1I) compared to control. However, there was an increase in the distance covered by the group OTC+FF when compared to DMSO group (F_(2, 84)_ = 4.434, p = 0.0142; Fig.1C). We observed an increase in the movement in OTC + FF compared to control and vehicle (H = 29.64, p < 0.0001; Fig. 1F).

**Figure 1.**
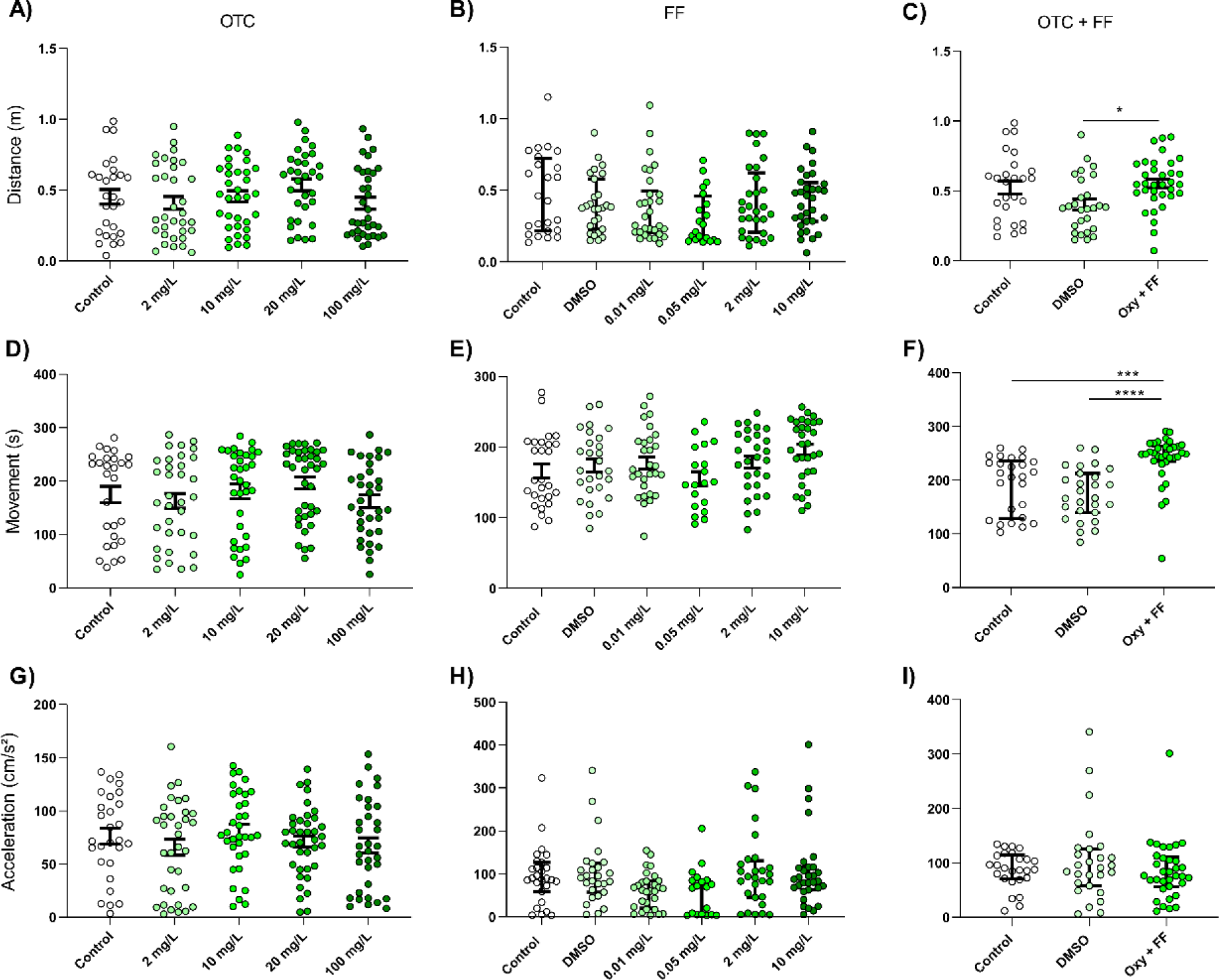
Effects of Oxytetracycline (OTC), Florfenicol (FF) or OTC+FF on exploratory behavior in zebrafish larvae. Distance covered, movement, and acceleration were evaluated for OTC (A, D, G), FF (B, E, H), and OTC + FF (C, F, I). Graphs A, C, E and G were analyzed using one-way analysis of variance (ANOVA) followed by Tukey test as a post hoc. Data from graphs are presented as mean ± SEM. Graphs B, D, F, H, and I were analyzed by Kruskal-Wallis following Dunn’s multiple comparisons test. Data from graphs are presented as median ± interquartile range (n = 20-34). * indicates significant difference at p ≤0.05, *** p = 0.0003, and **** p ≤0.0001. For statistical details see Results.

The turn angle did not alter to OTC- (H = 12.72, p = 0.0127; Fig. 2A) and FF-treated animals (H = 11.04, p = 0.0505; Fig. 2B). However, OTC + FF presented reduction of turn angle (H = 31.33, p < 0.0001; Fig. 2C) in relation to control and vehicle.

**Figure 2.**
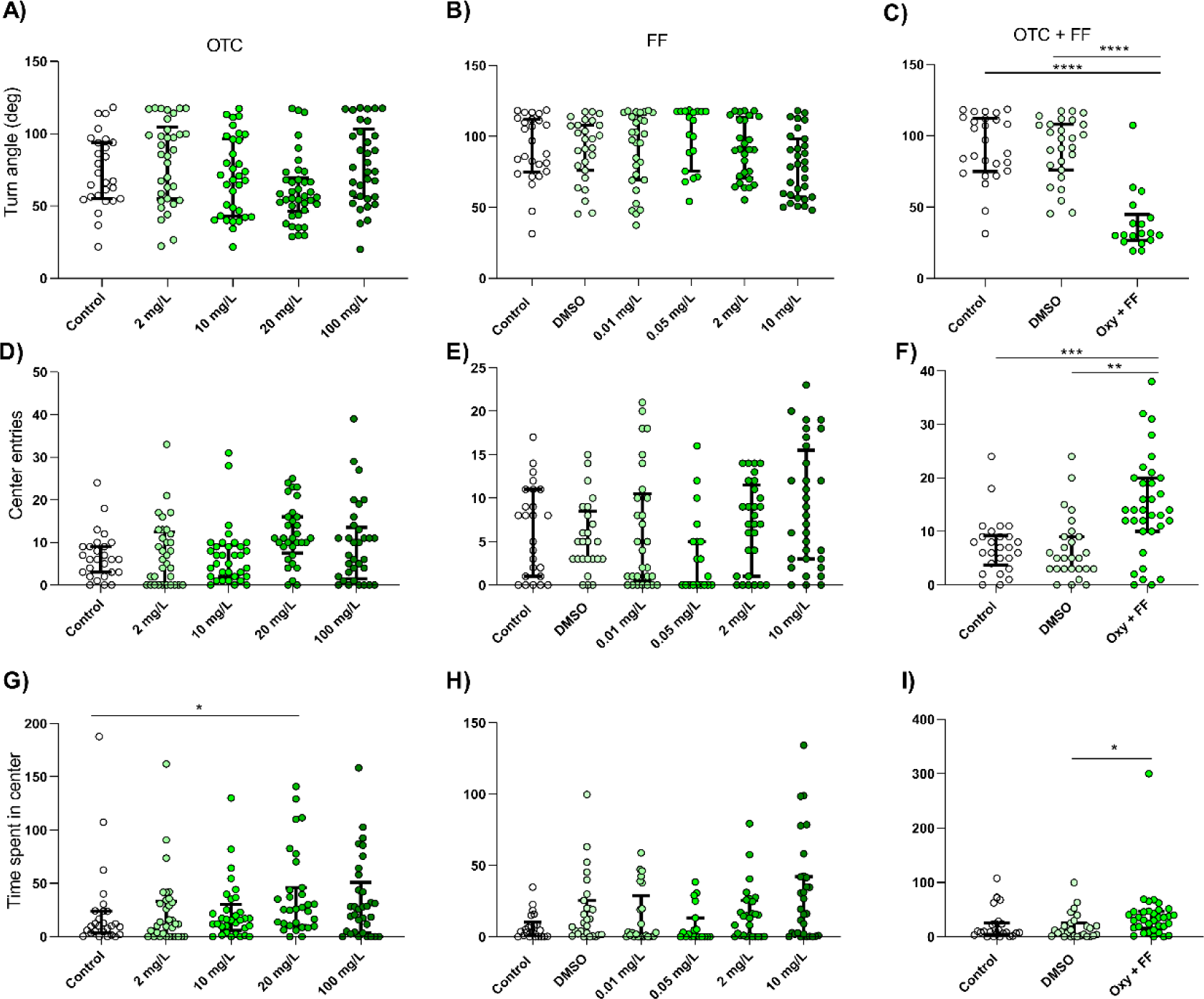
Effects of OTC, FF, and OTC+FF on anxiety-like behavior in zebrafish. Turn angle, Center entries, and time spent in center (s) were evaluated for OTC (A, D, G), FF (B, E, H), and OTC + FF (C, F, I). The data were analyzed by Kruskal-Wallis following Dunn’s multiple comparisons test. Data from graphs are presented as median ± interquartile range (n=20-34). * indicates significant difference at p ≤0.05, ** p = ≤0.5 and *** p ≤0.0005. For statistical details see Results.

For FF-treated animals, we did not observe differences in the number of entries in the center area (H = 11.07, p = 0.0500; Fig. 2E) and in the time spent in the center area (H = 13.75, p = 0.0172; Fig. 2H) when compared to control. For OTC-treated group, there were no differences in the number of center entries (H = 13.56, p = 0.0088; Fig. 2D), whereas an increase in the time spent in the center zone for animals treated with 20 mg/L OTC was observed (H = 9.696, p = 0.0459; Fig. 2G).

However, for OTC + FF-treated group, the number of entries compared to control or vehicle (H = 17.38, p = 0.0002; Fig. 2F) and time spent in the center (H = 9.179, p = 0.0102; Fig. 2I) were increased when compared to vehicle. The erratic movements did not demonstrate differences for OTC (H = 4.837, p = 0.3045, FF (F_(5, 154)_ = 2.011, p = 0.0801, and OTC + FF (H = 2.591, p = 0.2738; data not shown). The latencies to center entry to OTC (H = 3.045, p = 0.5504), FF (H = 7.043, p = 0.2174), and OTC + FF (H = 4.327, p = 0. 1149) groups did not show differences (data not shown).

### 3.2 Optomotor response

The optomotor response demonstrated a reduction after exposure to 10 mg/L and 20 mg/L OTC (F_(4, 130)_ = 7.921, p < 0.0001 Fig. 3A), as well as for OTC + FF (H = 19.56, p < 0.0001; Fig. 3C). Nevertheless, there were no differences from FF-treated group when compared to control (H = 10.59, p = 0.0601; Fig. 3B).

**Figure 3.**
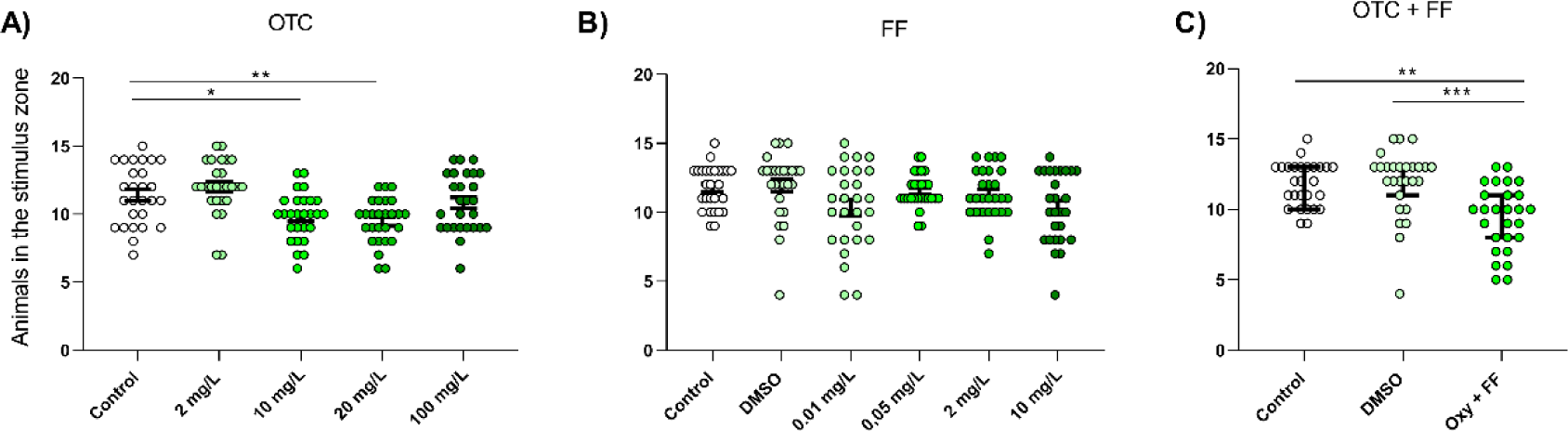
The optomotor response was evaluated in zebrafish larvae exposed to OTC (A), FF (B) or OTC + FF (C). Graph A was analyzed using one-way analysis of variance (ANOVA) followed by Tukey test as a post hoc, presented as mean ± SEM. Graphs B, and C were analyzed by Kruskal-Wallis following Dunn’s multiple comparisons test, presented as median ± interquartile range (n = 27). * indicates significant difference at p ≤ 0.05, ** = p ≤ 0.005, and *** p ≤ 0.0001. For statistical details see Results.

### 3.3 Aversive behavior

We did not observe any difference in aversive behavior for OTC- (F_(4, 38)_ = 1.193, p = 0.3298 Fig. 4A), FF- (F_(5, 48)_ = 0.9296, p = 0.4703 Fig. 4B), and OTC + FF-treated groups (F_(2, 24)_ = 0.1392, p = 0.8707 Fig. 4C).

**Figure 4.**
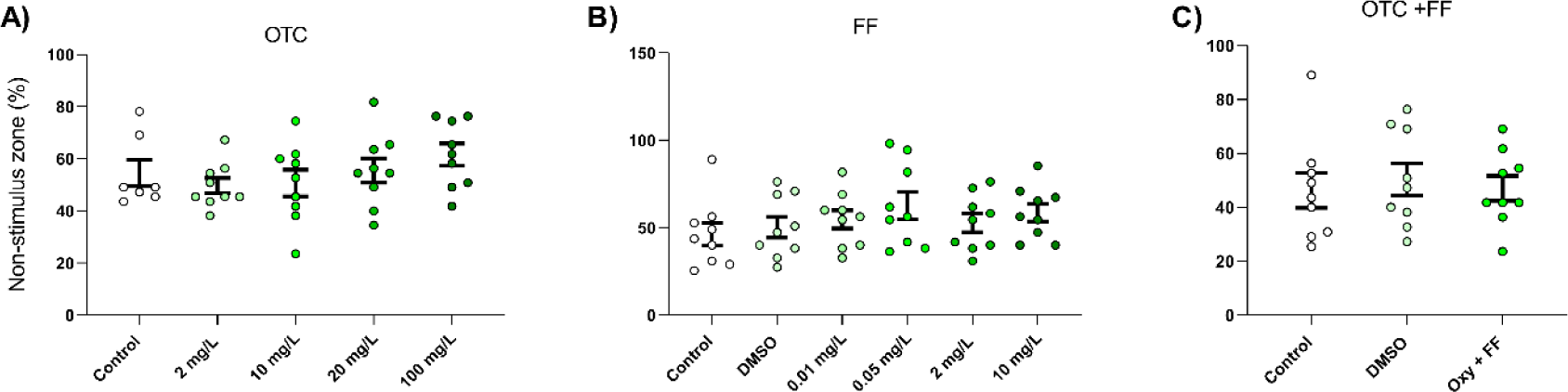
Effects of OTC (A), FF (B), and OTC + FF (C) on aversive stimulus in zebrafish larvae. The data were analyzed using one-way analysis of variance (ANOVA) followed by Tukey test as a post hoc. Data from graphs are presented as mean ± SEM (n = 9). For statistical details see Results.

## 4. Discussion

Zebrafish represent a sensitive organism to evaluate drug exposure through behavioral analysis. Here we described the behavioral effects caused by OTC or FF alone and both OTC + FF compounds.

The zebrafish exposed to OTC + FF showed specific alterations in pattern behavior. The movement was increased to OTC + FF exposure, but not for larvae exposed to OTC or FF alone. Elevated movement (mobile time) associated with higher distance covered, and acceleration have been considered a transient state of hyperactivity (Kalueff et al., 2013). Here we did not observe any difference in distance covered or acceleration to suggest hyperactivity. On the other hand, we observed larvae entered the central zone of the novel environment more often and stayed longer in the center area. When introduced in the novel environment, larvae are allowed to explore the arena to recognize and stay safe (Kysil et al., 2017; Stewart et al., 2014). Increased permanence in the central zone of an environment demonstrates less anxiety-like behavior. Studies have shown that zebrafish exposed to anxiolytic compounds stay longer in the center of the plate (Muniandy, 2018). Zebrafish with anxiety-like behavior traces remain longer in the peripheral part of the environment avoiding the center, coupled with a decrease in mobile time (Gusso et al., 2021; Wong et al., 2010). As prey, zebrafish try to defend themselves in the peripheral parts of the tank and darker places (Grossman et al., 2011; Gusso et al., 2021). However, as we observed, there is a reduced anxiety-like behavior in larvae exposed to OTC+ FF. In accordance, we observed that the turn angle, which represents the change in the direction of the animal through the center point (Abozaid et al., 2020; Nabinger et al., 2021) reduced in OTC + FF-treated animals. Studies suggested that the turn angle increased in response to anxiety-like behavior (Fulcher et al., 2017); however, our findings demonstrated that antibiotics reduced the turn angle in zebrafish larvae.

Our results present a somewhat different behavioral profile when compared to control zebrafish larvae. We observed an increased exploratory behavior through the distance traveled and the moving time, as well as a reduction in the turn angle. These results indicated a reduction of the anxious behavioral profile after antibiotics exposure in zebrafish larvae. We confirmed this when the OTC + FF-treated larvae spend less time following the lines in the optomotor stimulus compared to the control. The optomotor task has been designed to test responses to visual stimuli. When introduced into the test plate, the larvae tended to follow the black lines way. Conversely, the aversive stimulus that evaluates anxiety responses and the analysis of the escape behavior had no difference for any antibiotic tested. The aversive stimulus is commonly applied to avoidance tests and evades natural threats or predators (Nabinger et al., 2021; Pelkowski et al., 2011). The zebrafish larvae have visual perception enough to avoid the red ball (i.e., avoid a predator). However, they present a less anxious behavior than that described for 4dpf larvae, because they remained longer in the plate center, showed a turn angle reduction, and mobile time increased. Also, the larvae swan against the direction of optomotor stimuli. To perform sensorimotor behavior, the brain circuits need to be synchronous (Naumann et al., 2016). Once introduced and started the test, the zebrafish larvae usually follow the direction of the line. Changes in the optomotor test indicate impairment in visual, motor, or behavioral responses.

Antibiotics such as OTC and FF are commonly applied to prevent or treat diseases from several animal species, such as fish (Botelho et al., 2015; Leal et al., 2017). Each antibiotic presents different action mechanisms. OTC has been reported as an antibiotic that does not cross the blood-brain barrier. A study demonstrated high OTC levels in the fish intestine (Chemello et al., 2016). Studies have shown that OTC causes an imbalance in the gut microbiome while eliminating bacteria groups, allowing others to grow (Navarrete et al., 2008). In contrast, the FF penetration across the blood-brain barrier was 46% (Papich, 2016). The FF reduced not only the richness but also the diversity of the gut microbiota indicating an imbalance (Li et al., 2017). Here we have two compounds with different access to the brain. Neurotransmitter levels such as serotonin are regulated by the gut microbiome (Desbonnet et al., 2010) and most of these neurotransmitters come from the gut (Lu et al., 2019). The intestine has a crucial role in the brain through the gut-brain axis (Dinan and Cryan, 2017; Petra et al., 2015). Neurotransmitters of the intestine are loaded through the intestinal epithelium for the Vagus nerve and sensory nerves until the spinal cord to act in the brain (Slingerland and Stein-Thoeringer, 2018). The relationship of host-organism is harmonic; however, some pathological conditions have been described as associated with the gut-brain axis such as autism, depression, anxiety disorders, eating disorders, Parkinson’s disease, and pain syndromes (Slingerland and Stein-Thoeringer, 2018).

Interactions through the gut recognition mechanism result in numerous functional networks that are yet to be clarified. The variability of the microbiota is related to chronic diseases (Clavel and Ecker, 2018). On the other hand, during a transitional period, antibiotics can alter behaviors, such as anxiety-like behavior (Han and Kim, 2019; Petra et al., 2015). The behavioral analyses can provide insights into how antibiotics may be linked to the altered function of the central nervous system. A broad range of chemical compounds could cause brain dysfunction, even with no apparent effects. Studies need to be conducted to better understand the effects of the microbiome on the brain after antibiotic exposure and their impacts on behavioral responses.

## 5. Conclusion

In summary, this study demonstrated behavioral responses induced by OTC + FF that reduced anxiety-like behavior. This study reinforces the importance of investigating the effect of antibiotics on behavioral responses and their mechanisms since these compounds are widely used in fish production.

## Acknowledgments

This study was financed in part by the Coordenação de Aperfeiçoamento de Pessoal de Nível Superior - Brasil (CAPES) - finance code 001, Conselho Nacional de Desenvolvimento Científico e Tecnológico (CNPq; Proc. 420695/2018-4), Fundação de Amparo à Pesquisa do Estado do Rio Grande do Sul (FAPERGS; 17/2551-0000977-0) and Instituto Nacional de Ciências e Tecnologia para Doenças Cerebrais, Excitotoxicidade e Neuroproteção. C.D.B. (Proc. 304450/2019-7) was the recipients of a fellowship from CNPq.

## References

Abozaid, A., Tsang, B., Gerlai, R., 2020. The effects of small but abrupt change in temperature on the behavior of larval zebrafish. Physiol. Behav. 227, 113169. https://doi.org/10.1016/j.physbeh.2020.113169

Almeida, A.R., Tacão, M., Machado, A.L., Golovko, O., Zlabek, V., Domingues, I., Henriques, I., 2019. Long-term effects of oxytetracycline exposure in zebrafish: A multi-level perspective. Chemosphere 222, 333–344. https://doi.org/10.1016/j.chemosphere.2019.01.147

Botelho, R.G., Christofoletti, C.A., Correia, J.E., Ansoar, Y., Olinda, R.A., Tornisielo, V.L., 2015. Genotoxic responses of juvenile tilapia (Oreochromis niloticus) exposed to florfenicol and oxytetracycline. Chemosphere 132, 206–212. https://doi.org/10.1016/j.chemosphere.2015.02.053

Bridi, D., Altenhofen, S., Gonzalez, J.B., Reolon, G.K., Bonan, C.D., 2017. Glyphosate and Roundup ® alter morphology and behavior in zebrafish. Toxicology 392, 32–39. https://doi.org/10.1016/j.tox.2017.10.007

Cachat, J., Stewart, A., Grossman, L., Gaikwad, S., Kadri, F., Chung, K.M., Wu, N., Wong, K., Roy, S., Suciu, C., Goodspeed, J., Elegante, M., Bartels, B., Elkhayat, S., Tien, D., Tan, J., Denmark, A., Gilder, T., Kyzar, E., Dileo, J., Frank, K., Chang, K., Utterback, E., Hart, P., Kalueff, A. V., 2010. Measuring behavioral and endocrine responses to novelty stress in adult zebrafish. Nat. Protoc. 5, 1786–1799. https://doi.org/10.1038/nprot.2010.140

Chemello, G., Piccinetti, C., Randazzo, B., Carnevali, O., Maradonna, F., Magro, M., Bonaiuto, E., Vianello, F., Radaelli, G., Fifi, A.P., Gigliotti, F., Olivotto, I., 2016. Oxytetracycline Delivery in Adult Female Zebrafish by Iron Oxide Nanoparticles. Zebrafish 13, 495–503. https://doi.org/10.1089/zeb.2016.1302

Clavel, T., Ecker, J., 2018. Microbiome and Diseases: Metabolic Disorders, in: The Gut Microbiome in Health and Disease. Springer International Publishing, Cham, pp. 251–277. https://doi.org/10.1007/978-3-319-90545-7_16

Colwill, R.M., Creton, R., 2011. Locomotor behaviors in zebrafish (Danio rerio) larvae. Behav. Processes 86, 222–229. https://doi.org/10.1016/j.beproc.2010.12.003

Desbonnet, L., Garrett, L., Clarke, G., Kiely, B., Cryan, J.F., Dinan, T.G., 2010. Effects of the probiotic Bifidobacterium infantis in the maternal separation model of depression. Neuroscience 170, 1179–1188. https://doi.org/10.1016/j.neuroscience.2010.08.005

Dinan, T.G., Cryan, J.F., 2017. Brain-Gut-Microbiota Axis and Mental Health. Psychosom. Med. 79, 920–926. https://doi.org/10.1097/PSY.0000000000000519

EMA - European Medicines Agency, 2020. Advice on implementing measures under Article 106 (6) of Regulation (EU) 2019/6 on veterinary medicinal products – scientific problem analysis and recommendations to ensure a safe and efficient administration of oral veterinary medicinal products via rout [WWW Document]. URL https://www.ema.europa.eu/en/documents/regulatory-procedural-guideline/advice-implementing-measures-under-article-106-6-regulation-eu-2019/6-veterinary-medicinal-products-scientific-problem-analysis-recommendations-ensure-safe-efficient_en.pdf (accessed 4.6.21).

FAO, 2020. The State of World Fisheries and Aquaculture 2020 [WWW Document]. URL http://www.fao.org/3/ca9229en/CA9229EN.pdf (accessed 3.31.21).

FDA - Food and Drug Administration, 2021. Approved Aquaculture Drugs [WWW Document]. URL https://www.fda.gov/animal-veterinary/aquaculture/approved-aquaculture-drugs (accessed 4.6.21).

Fulcher, N., Tran, S., Shams, S., Chatterjee, D., Gerlai, R., 2017. Neurochemical and Behavioral Responses to Unpredictable Chronic Mild Stress Following Developmental Isolation: The Zebrafish as a Model for Major Depression. Zebrafish 14, 23–34. https://doi.org/10.1089/zeb.2016.1295

Gaspary, K.V., Reolon, G.K., Gusso, D., Bonan, C.D., 2018. Novel object recognition and object location tasks in zebrafish: Influence of habituation and NMDA receptor antagonism. Neurobiol. Learn. Mem. 155, 249–260. https://doi.org/10.1016/j.nlm.2018.08.005

Gerlai, R., 2016. Learning and memory in zebrafish (Danio rerio). pp. 551–586. https://doi.org/10.1016/bs.mcb.2016.02.005

Gerlai, R., 2003. Zebrafish: An uncharted behavior genetic model. Behav. Genet. 33, 1–8. https://doi.org/10.2144/000113931

Gerlai, R., Lahav, M., Guo, S., Rosenthal, A., 2000. Drinks like a fish: Zebra fish (Danio rerio) as a behavior genetic model to study alcohol effects. Pharmacol. Biochem. Behav. 67, 773–782. https://doi.org/10.1016/S0091-3057(00)00422-6

Giacomini, A.C.V.V., de Abreu, M.S., Koakoski, G., Idalêncio, R., Kalichak, F., Oliveira, T.A., da Rosa, J.G.S., Gusso, D., Piato, A.L., Barcellos, L.J.G., 2015. My stress, our stress: Blunted cortisol response to stress in isolated housed zebrafish. Physiol. Behav. 139. https://doi.org/10.1016/j.physbeh.2014.11.035

Grossman, L., Stewart, A., Gaikwad, S., Utterback, E., Wu, N., DiLeo, J., Frank, K., Hart, P., Howard, H., Kalueff, A. V., 2011. Effects of piracetam on behavior and memory in adult zebrafish. Brain Res. Bull. 85, 58–63. https://doi.org/10.1016/j.brainresbull.2011.02.008

Gusso, D., Altenhofen, S., Fritsch, P.M., Rübensam, G., Bonan, C.D., 2021. Oxytetracycline induces anxiety-like behavior in adult zebrafish. Toxicol. Appl. Pharmacol. 426, 115616. https://doi.org/10.1016/j.taap.2021.115616

Gusso, D., Reolon, G.K., Gonzalez, J.B., Altenhofen, S., Kist, L.W., Bogo, M.R., Bonan, C.D., 2020. Pyriproxyfen Exposure Impairs Cognitive Parameters and Alters Cortisol Levels in Zebrafish. Front. Behav. Neurosci. 14, 32–39. https://doi.org/10.3389/fnbeh.2020.00103

Hallare, A., Nagel, K., Köhler, H.-R., Triebskorn, R., 2006. Comparative embryotoxicity and proteotoxicity of three carrier solvents to zebrafish (Danio rerio) embryos. Ecotoxicol. Environ. Saf. 63, 378–388. https://doi.org/10.1016/j.ecoenv.2005.07.006

Han, S.-K., Kim, D.H., 2019. Lactobacillus mucosae and Bifidobacterium longum Synergistically Alleviate Immobilization Stress-Induced Anxiety/Depression in Mice by Suppressing Gut Dysbiosis. J. Microbiol. Biotechnol. 29, 1369–1374. https://doi.org/10.4014/jmb.1907.07044

Harnisz, M., Korzeniewska, E., Gołas, I., 2015. The impact of a freshwater fish farm on the community of tetracycline-resistant bacteria and the structure of tetracycline resistance genes in river water. Chemosphere 128, 134–141. https://doi.org/10.1016/j.chemosphere.2015.01.035

Henriksson, P.J.G., Rico, A., Troell, M., Klinger, D.H., Buschmann, A.H., Saksida, S., Chadag, M. V., Zhang, W., 2018. Unpacking factors influencing antimicrobial use in global aquaculture and their implication for management: a review from a systems perspective. Sustain. Sci. 13, 1105–1120. https://doi.org/10.1007/s11625-017-0511-8

Idalencio, R., Kalichak, F., Rosa, J.G.S., Oliveira, T.A. de, Koakoski, G., Gusso, D., Abreu, M.S. de, Giacomini, A.C.V., Barcellos, H.H. de A., Piato, A.L., Barcellos, L.J.G., 2015. Waterborne Risperidone Decreases Stress Response in Zebrafish. PLoS One 10, e0140800. https://doi.org/10.1371/journal.pone.0140800

Jang, H.-M., Lee, H.-J., Jang, S.-E., Han, M.J., Kim, D.-H., 2018. Evidence for interplay among antibacterial-induced gut microbiota disturbance, neuro-inflammation, and anxiety in mice. Mucosal Immunol. 11, 1386–1397. https://doi.org/10.1038/s41385-018-0042-3

Kalueff, A. V, Gebhardt, M., Stewart, A.M., Cachat, J.M., Brimmer, M., Chawla, J.S., Craddock, C., Kyzar, E.J., Roth, A., Landsman, S., Gaikwad, S., Robinson, K., Baatrup, E., Tierney, K., Shamchuk, A., Norton, W., Miller, N., Nicolson, T., Braubach, O., Gilman, C.P., Pittman, J., Rosemberg, D.B., Gerlai, R., Echevarria, D., Lamb, E., Neuhauss, S.C.F., 2013. Towards a Comprehensive Catalog of Zebrafish 10, 70–86. https://doi.org/10.1089/zeb.2012.0861

Kayani, M. ur R., Yu, K., Qiu, Y., Shen, Y., Gao, C., Feng, R., Zeng, X., Wang, W., Chen, L., Su, H.L., 2021. Environmental concentrations of antibiotics alter the zebrafish gut microbiome structure and potential functions. Environ. Pollut. 22, 116760. https://doi.org/10.1016/j.envpol.2021.116760

Kysil, E. V., Meshalkina, D.A., Frick, E.E., Echevarria, D.J., Rosemberg, D.B., Maximino, C., Lima, M.G., Abreu, M.S., Giacomini, A.C., Barcellos, L.J.G., Song, C., Kalueff, A. V., 2017. Comparative Analyses of Zebrafish Anxiety-Like Behavior Using Conflict-Based Novelty Tests. Zebrafish 14, 197–208. https://doi.org/10.1089/zeb.2016.1415

Laxminarayan, R., Duse, A., Wattal, C., Zaidi, A.K.M., Wertheim, H.F.L., Sumpradit, N., Vlieghe, E., Hara, G.L., Gould, I.M., Goossens, H., Greko, C., So, A.D., Bigdeli, M., Tomson, G., Woodhouse, W., Ombaka, E., Peralta, A.Q., Qamar, F.N., Mir, F., Kariuki, S., Bhutta, Z.A., Coates, A., Bergstrom, R., Wright, G.D., Brown, E.D., Cars, O., 2013. Antibiotic resistance—the need for global solutions. Lancet Infect. Dis. 13, 1057–1098. https://doi.org/10.1016/S1473-3099(13)70318-9

Leal, J.F., Henriques, I.S., Correia, A., Santos, E.B.H., Esteves, V.I., 2017. Antibacterial activity of oxytetracycline photoproducts in marine aquaculture’s water. Environ. Pollut. 220, 644–649. https://doi.org/10.1016/j.envpol.2016.10.021

Li, J., Dong, T., Keerthisinghe, T.P., Chen, H., Li, M., Chu, W., Yang, J., Hu, Z., Allen, S., Dong, W., Fang, M., 2020. Long-term oxytetracycline exposure potentially alters brain thyroid hormone and serotonin homeostasis in zebrafish. J. Hazard. Mater. 399, 123061. https://doi.org/10.1016/j.jhazmat.2020.123061

Li, R., Wang, H., Shi, Q., Wang, N., Zhang, Z., Xiong, C., Liu, J., Chen, Y., Jiang, L., Jiang, Q., 2017. Effects of oral florfenicol and azithromycin on gut microbiota and adipogenesis in mice. PLoS One 12, e0181690. https://doi.org/10.1371/journal.pone.0181690

Lu, Y., Zhang, Z., Liang, X., Chen, Y., Zhang, J., Yi, H., Liu, T., Yang, L., Shi, H., Zhang, L., 2019. Study of gastrointestinal tract viability and motility via modulation of serotonin in a zebrafish model by probiotics. Food Funct. 10, 7416–7425. https://doi.org/10.1039/C9FO02129A

Lurie, I., Yang, Y.-X., Haynes, K., Mamtani, R., Boursi, B., 2015. Antibiotic Exposure and the Risk for Depression, Anxiety, or Psychosis. J. Clin. Psychiatry 76, 1522–1528. https://doi.org/10.4088/JCP.15m09961

Miller, N.Y., Gerlai, R., 2008. Oscillations in shoal cohesion in zebrafish (Danio rerio). Behav. Brain Res. 193, 148–151. https://doi.org/10.1016/j.bbr.2008.05.004

Mo, W.Y., Chen, Z., Leung, H.M., Leung, A.O.W., 2017. Application of veterinary antibiotics in China’s aquaculture industry and their potential human health risks. Environ. Sci. Pollut. Res. 24, 8978–8989. https://doi.org/10.1007/s11356-015-5607-z

Monteiro, S.H., Garcia, F., Gozi, K.S., Romera, D.M., Francisco, J.G., Moura-Andrade, G.C.R., Tornisielo, V.L., 2016. Relationship between antibiotic residues and occurrence of resistant bacteria in Nile tilapia (Oreochromisniloticus) cultured in cage-farm. J. Environ. Sci. Heal. Part B 51, 817–823. https://doi.org/10.1080/03601234.2016.1208457

Müller, T.E., Fontana, B.D., Bertoncello, K.T., Franscescon, F., Mezzomo, N.J., Canzian, J., Stefanello, F. V., Parker, M.O., Gerlai, R., Rosemberg, D.B., 2020. Understanding the neurobiological effects of drug abuse: Lessons from zebrafish models. Prog. Neuro-Psychopharmacology Biol. Psychiatry 100, 109873. https://doi.org/10.1016/j.pnpbp.2020.109873

Muniandy, Y., 2018. The Use of Larval Zebrafish (Danio rerio) Model for Identifying New Anxiolytic Drugs from Herbal Medicine. Zebrafish 15, 321–339. https://doi.org/10.1089/zeb.2018.1562

Nabinger, D.D., Altenhofen, S., Bitencourt, P.E.R., Nery, L.R., Leite, C.E., Vianna, M.R.M.R., Bonan, C.D., 2018. Nickel exposure alters behavioral parameters in larval and adult zebrafish. Sci. Total Environ. 624, 1623–1633. https://doi.org/10.1016/j.scitotenv.2017.10.057

Nabinger, D.D., Altenhofen, S., Peixoto, J.V., da Silva, J.M.K., Gerlai, R., Bonan, C.D., 2021. Feeding status alters exploratory and anxiety-like behaviors in zebrafish larvae exposed to quinpirole. Prog. Neuro-Psychopharmacology Biol. Psychiatry 108, 110179. https://doi.org/10.1016/j.pnpbp.2020.110179

Naumann, E.A., Fitzgerald, J.E., Dunn, T.W., Rihel, J., Sompolinsky, H., Engert, F., 2016. From Whole-Brain Data to Functional Circuit Models: The Zebrafish Optomotor Response. Cell 167, 947–960.e20. https://doi.org/10.1016/j.cell.2016.10.019

Navarrete, P., Mardones, P., Opazo, R., Espejo, R., Romero, J., 2008. Oxytetracycline Treatment Reduces Bacterial Diversity of Intestinal Microbiota of Atlantic Salmon. J. Aquat. Anim. Health 20, 177–183. https://doi.org/10.1577/H07-043.1

Naylor, R.L., Hardy, R.W., Buschmann, A.H., Bush, S.R., Cao, L., Klinger, D.H., Little, D.C., Lubchenco, J., Shumway, S.E., Troell, M., 2021. A 20-year retrospective review of global aquaculture. Nat. | 591, 551.

Nery, L.R., Eltz, N.S., Hackman, C., Fonseca, R., Altenhofen, S., Guerra, H.N., Freitas, V.M., Bonan, C.D., Vianna, M.R.M.R., 2014. Brain intraventricular injection of amyloid-β in zebrafish embryo impairs cognition and increases tau phosphorylation, effects reversed by lithium. PLoS One 9, e105862. https://doi.org/10.1371/journal.pone.0105862

ONU, 2019. World population prospects 2019, Department of Economic and Social Affairs. World Population Prospects 2019.

Papich, M.G., 2016. Florfenicol, in: Saunders Handbook of Veterinary Drugs. Elsevier, pp. 327–329. https://doi.org/10.1016/B978-0-323-24485-5.00264-3

Pelkowski, S.D., Kapoor, M., Richendrfer, H.A., Wang, X., Colwill, R.M., Creton, R., 2011. A novel high-throughput imaging system for automated analyses of avoidance behavior in zebrafish larvae. Behav. Brain Res. 223, 135–144. https://doi.org/10.1016/j.bbr.2011.04.033

Petersen, B.D., Pereira, T.C.B., Altenhofen, S., Nabinger, D.D., Ferreira, P.M. de A., Bogo, M.R., Bonan, C.D., 2021. Antibiotic drugs alter zebrafish behavior. Comp. Biochem. Physiol. Part C Toxicol. Pharmacol. 242, 108936. https://doi.org/10.1016/j.cbpc.2020.108936

Petra, A.I., Panagiotidou, S., Hatziagelaki, E., Stewart, J.M., Conti, P., Theoharides, T.C., 2015. Gut-Microbiota-Brain Axis and Its Effect on Neuropsychiatric Disorders with Suspected Immune Dysregulation. Clin. Ther. https://doi.org/10.1016/j.clinthera.2015.04.002

Rok, J., Wrześniok, D., Beberok, A., Otręba, M., Delijewski, M., Buszman, E., 2017. Phototoxic effect of oxytetracycline on normal human melanocytes. Toxicol. Vitr. 48, 26–32. https://doi.org/10.1016/j.tiv.2017.12.008

Saito-Shida, S., Shiono, K., Narushima, J., Nemoto, S., Akiyama, H., 2019. Determination of total florfenicol residues as florfenicol amine in bovine tissues and eel by liquid chromatography–tandem mass spectrometry using external calibration. J. Chromatogr. B 1109, 37–44. https://doi.org/10.1016/j.jchromb.2019.01.018

San Martín, B., Fresno, M., Cornejo, J., Godoy, M., Ibarra, R., Vidal, R., Araneda, M., Anadón, A., Lapierre, L., 2019. Optimization of florfenicol dose against Piscirickettsia salmonis in Salmo salar through PK/PD studies. PLoS One 14, e0215174. https://doi.org/10.1371/journal.pone.0215174

Schar, D., Klein, E.Y., Laxminarayan, R., Gilbert, M., Van Boeckel, T.P., 2020. Global trends in antimicrobial use in aquaculture. Sci. Rep. 10, 21878. https://doi.org/10.1038/s41598-020-78849-3

Slingerland, A.E., Stein-Thoeringer, C.K., 2018. Microbiome and Diseases: Neurological Disorders, in: The Gut Microbiome in Health and Disease. Springer International Publishing, Cham, pp. 295–310. https://doi.org/10.1007/978-3-319-90545-7_18

Stewart, A.M., Braubach, O., Spitsbergen, J., Gerlai, R., Kalueff, A. V., 2014. Zebrafish models for translational neuroscience research: from tank to bedside. Trends Neurosci. 37, 264–278. https://doi.org/10.1016/j.tins.2014.02.011

Wernli, D., Jørgensen, P.S., Parmley, E.J., Troell, M., Majowicz, S., Harbarth, S., Léger, A., Lambraki, I., Graells, T., Henriksson, P.J.G., Carson, C., Cousins, M., Skoog Ståhlgren, G., Mohan, C. V., Simpson, A.J.H., Wieland, B., Pedersen, K., Schneider, A., Chandy, S.J., Wijayathilaka, T.P., Delamare-Deboutteville, J., Vila, J., Stålsby Lundborg, C., Pittet, D., 2020. Evidence for action: a One Health learning platform on interventions to tackle antimicrobial resistance. Lancet Infect. Dis. 20, e307–e311. https://doi.org/10.1016/S1473-3099(20)30392-3

Westerfield, M., 2007. The Zebrafish Book. A Guide for the Laboratory Use of Zebrafish (Danio rerio), 5th Edition. Univ. Oregon Press. Eugene.

Wong, K., Elegante, M., Bartels, B., Elkhayat, S., Tien, D., Roy, S., Goodspeed, J., Suciu, C., Tan, J., Grimes, C., Chung, A., Rosenberg, M., Gaikwad, S., Denmark, A., Jackson, A., Kadri, F., Chung, K.M., Stewart, A., Gilder, T., Beeson, E., Zapolsky, I., Wu, N., Cachat, J., Kalueff, A. V., 2010. Analyzing habituation responses to novelty in zebrafish (Danio rerio). Behav. Brain Res. 208, 450–457. https://doi.org/10.1016/j.bbr.2009.12.023

Yu, K., Li, X., Qiu, Y., Zeng, X., Yu, X., Wang, W., Yi, X., Huang, L., 2020. Low-dose effects on thyroid disruption in zebrafish by long-term exposure to oxytetracycline. Aquat. Toxicol. 227, 105608. https://doi.org/10.1016/j.aquatox.2020.105608

